# Aeration Strategy at Birth Does Not Impact Carotid Haemodynamics in Preterm Lambs

**DOI:** 10.1101/2022.03.03.482754

**Authors:** Sophia I Dahm, Kelly R Kenna, David Stewart, Prudence M Pereira-Fantini, Karen E McCall, Liz Perkins, Magdy Sourial, David G Tingay

## Abstract

**Background:** The impact of different respiratory strategies at birth on the preterm lung is well understood, however, concerns have been raised that lung recruitment may impede cerebral haemodynamics. This study aims to examine the effect of three different ventilation strategies on cerebral haemodynamics and oxygenation.

**Methods:** 124-127 day gestation apnoeic intubated preterm lambs (n=68) being studied as part of a larger program primarily assessing lung injury were randomised to positive pressure ventilation with positive end-expiratory pressure (PEEP) 8 cmH_2_O (No-RM; n=12), sustained inflation (SI; n=15) or dynamic PEEP strategy (DynPEEP; maximum PEEP 14 or 20 cmH_2_O, n=41) at birth, followed by 90 minutes of standardised ventilation. Haemodynamic data was continuously recorded, with intermittent arterial blood gas analysis. Main outcome measures for this study included carotid blood flow, carotid artery oxygen content and carotid oxygen delivery.

**Results:** Overall carotid blood flow measures were comparable between strategies, with the exception of mean carotid blood flow which was significantly lower for the SI group compared to the No-RM and DynPEEP groups respectively over the first 3 minutes (p<0.0001, mixed-effects model). Carotid oxygen content and oxygen delivery were similar between strategies. Maximum PEEP did not alter cerebral haemodynamic measures.

**Conclusion:** Although there were some short-term variations in cerebral haemodynamics between different PEEP strategies and SI, these were not sustained.

**Impact:**

- Different pressure strategies to facilitate lung aeration at birth in preterm infants have been proposed. There is minimal information on the effect of lung recruitment on cerebral haemodynamics.
- This is the first study that compares the effect of sustained lung inflation, and dynamic and static positive end-expiratory pressure on cerebral haemodynamics.
- We found that the different ventilation strategies did not alter carotid blood flow, carotid oxygen content or carotid oxygen delivery.
- This preclinical study provides some reassurance that respiratory strategies designed to focus on lung aeration at birth may not impact cerebral haemodynamics in preterm neonates.

## INTRODUCTION

Lung aeration is a fundamental component of the transition to air-breathing at birth.^1^ Achieving and maintaining lung aeration is challenging for many preterm infants and there has been considerable interest in the optimal methods of supporting lung aeration in preterm infants, including the use of adequate positive end-expiratory pressure (PEEP) or an initial sustained inflation (SI).^2–6^ Preclinical studies have repeatedly demonstrated that the use of adequate PEEP at birth, and preferably dynamic PEEP manoeuvres, optimises lung mechanics, aeration and reduces the risks of lung injury compared to SI.^2–6^ Recent clinical trials of fixed-duration SI in preterm infants have suggested potentially important adverse outcomes without lung protective benefits. Studies assessing ventilation strategies have primarily focused on respiratory outcomes. To date, there have been minimal reports detailing cerebrovascular responses to PEEP approaches at birth.

Cerebral blood flow (CeBF) decreases after birth in both healthy term lambs and infants as a proposed response to increasing oxygen saturation and cerebral oxygen delivery following spontaneous breathing.^8,9^ However, the behaviour of CeBF and oxygen delivery in preterm infants aerated by PEEP recruitment or SI at birth remains largely unknown. Whilst its use is ubiquitous, PEEP strategies at birth have not been subjected to systematic evaluation in preterm infants, resulting in the use of variable PEEP levels.^10^ A SI at birth increased CeBF and carotid oxygen content compared to a low static PEEP with tidal inflations in preterm lambs.^11^ Beyond the respiratory transition at birth, excessive PEEP levels impacts haemodynamics in preterm lambs.^12,13^ The impact of higher PEEP levels at birth is less clear, with a recent pilot lamb study suggesting engorgement of micro-vessels and an increase in CeBF.^14^

We hypothesised that measures of cerebral haemodynamics will be impacted by different PEEP levels or a SI at birth. We aimed to determine the effect of static and dynamic PEEP level strategies or a SI to facilitate lung aeration at birth on carotid blood flow (CBF), carotid arterial blood oxygen (CAO) content and carotid oxygen delivery in intubated preterm lambs.

## METHODS

### Protocol

All techniques and procedures were approved by the Animal Ethics Committee of the Murdoch Children’s Research Institute (MCRI), Melbourne, Australia in accordance with National Health and Medical Research Council (NHMRC) guidelines. This study was performed as part of a large inter-linked preterm lamb study program that aimed to determine the optimal methods of applying positive pressure ventilation (PPV) at birth. In the interest of animal reduction, and with the prior knowledge of our Animal Ethics Committee and funding body (NHMRC), some ventilation groups have been reported in different studies with distinctly separate aims.^2–4^

### Subjects

The detailed methodology and complete set of outcome measures for the broader study has been reported previously.^2–5^ In summary, 124-127d, preterm lambs born via Caesarean section under general anaesthesia to betamethasone-treated (11.7mg) ewes were instrumented before delivery. All ewes were from the same flock and studied over two breeding seasons of similar environmental factors. Instrumentation was performed on exteriorisation of the fetal chest, and included endotracheal tube intubation (4.0 mm cuffed), passive drainage of fetal lung fluid, flow probe placement around a carotid artery (3 mm Transonic, AD Instruments, Sydney, Australia) and occlusive cannulation of the contralateral carotid artery and external jugular vein. At delivery, the umbilical cord was cut and the lamb moved to a resuscitaire and weighed prior to commencing respiratory support (maximum delay between cord cut and first inflation 1 minute). CBF, ventilation data, peripheral oxygen saturation (SpO_2_), heart rate and arterial blood pressure were recorded throughout (200 Hz, LabChart V7, AD Instruments, Sydney, Australia). Arterial blood gas (ABG) analysis was performed at fetal (Cord Intact; CI), 5, 15 minutes and then 15 minutely after birth. Ketamine and midazolam infusions were titrated to minimum doses that suppressed breathing and maintained analgesia.

### Measurements

As described in detail previously,^2,4^ lambs were randomised to either **1)** PPV at a PEEP 8 cmH_2_O with volume targeted ventilation (VTV) from birth for 90 min (**No-RM**), **2)** an initial **SI** delivered at 35 cmH_2_O until aeration stability was achieved on real-time imaging of lung volume using electrical impedance tomography (mean 153.2 (24.3) seconds),^2,4,6^ or **3)** a dynamic PEEP manoeuvre delivered over 3 min using PEEP steps of 2 cmH_2_O to a maximum of 14 cmH_2_O (20s increments/PEEP step) or 20 cmH_2_O (10s increments), depending on group allocation, followed by the same stepwise decreases to a final PEEP of 8 cmH_2_O (**DynPEEP**). After the initial intervention, the SI and DynPEEP groups were managed as per No-RM. Fraction of inspired oxygen and tidal volume (V_T_) were started at 0.3 and 7 ml/kg respectively, and both titrated using a standardised strategy to maintain pre-ductal SpO_2_ 88-94% and arterial carbon dioxide 45-60 mmHg following the 5 minute ABG, with surfactant therapy at 10 minutes.^2^

A two minute measurement of CBF and cardiorespiratory data were made immediately prior to cord clamping. CBF data were extracted at 10s intervals during the fetal period, both intact cord (CI) and after cord clamping (CC) prior to commencing ventilation. CBF data were then extracted over 10s at minutely intervals to 5 minutes after starting respiratory support, and over 20s intervals at 10, 15, 30 and 90 minutes. In the DynPEEP groups, measures were made at maximum PEEP (80s) and end of dynamic PEEP (8 cmH_2_O; 140s) rather than 1 and 2 minutes. In all groups, measures from 3 to 90 minutes were at PEEP 8 cmH_2_O. From the CBF data, the mean waveform amplitude, maximum, minimum, and overall mean were extracted. Carotid blood flow is non-linear with dependency on heart rate and systemic blood pressure, complicated by the possibility of the presence of a ductus arteriosus in a fetus. Multiple waveform measurements were, therefore, extracted to provide a comprehensive analysis of carotid blood flow. The maximum CBF corresponds to systolic blood pressure values, whilst the minimum corresponds to diastolic values. The amplitude measures the difference between the systolic and diastolic flow values (height of the waveform), whilst the mean incorporates all phases of the carotid blood flow. Values were then expressed by birth weight. In this study, the contralateral carotid artery was occluded for vascular access and, therefore, the absolute CBF refers to the flow measured in the remaining patent carotid artery. As occlusion of one carotid artery may influence the pattern of upper body flow distribution, we presented the data as both change from baseline for accuracy and in absolute values for ease of understanding. CAO content and carotid oxygen delivery were calculated from the ABG results using the following formulas:

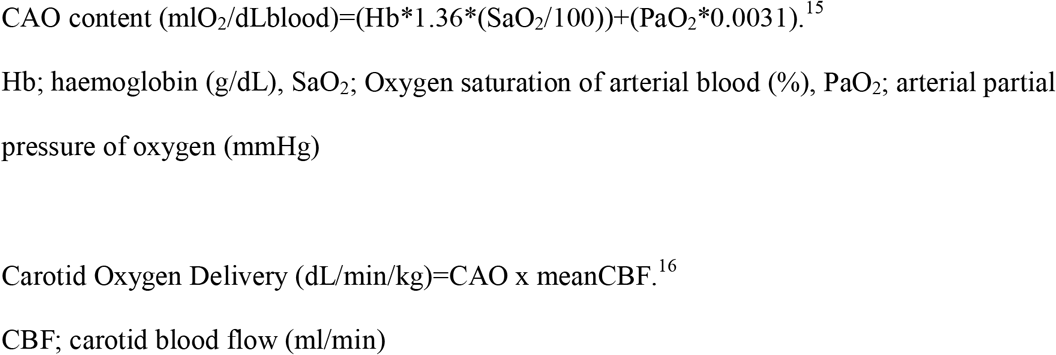

### Data analysis

Sample size was determined by the primary studies which were powered to lung injury primary outcomes, and required larger group sizes (approximately 20 lambs/group) than studies powered to cardiorespiratory measures.^2^ Previous preterm lamb studies reporting CBF and CAO content have shown important differences with 6/group.^11,17^. Our broader study was designed to investigate different ventilation strategies at birth, specifically SI and DynPEEP. In the interest of reduction, the No-RM group acted as a control across multiple studies. This resulted in an unbalanced number of lambs in each group. As reported in the original studies, there was no difference in adverse event rates between groups. Data were analysed using a mixed-effect model (Tukey post-tests) using ventilation strategy and time as principle variables. Within the DynPEEP group, subgroup analysis comparing the maximum delivered PEEP (14 vs. 20 cmH_2_O) was performed using a two-way ANOVA. Analysis was performed in PRISM V9 (GraphPad, San Diego, CA) and p<0.05 considered significant.

## RESULTS

A total of 89 lambs from our data set were available for analysis. 21 lambs were excluded because of unreliable CBF measures related to probe-vessel contact. No lambs were excluded based on clinical stability. Table 1 and Online Supplement Table 1 describe the clinical characteristics of the 68 lambs included for analysis. Incomplete ABG data in 6 lambs resulted in 62 lambs available for calculation of CAO content and carotid oxygen delivery. The lambs were well matched. There were statistical but not clinically significant differences in the amount of fetal lung fluid drained before birth (No-RM higher), fetal PaO_2_ (No-RM lower) and minimum CBF and PaCO_2_ (both SI higher), with no significant differences in the absolute fetal CI mean and maximum CBF values.

Figure 1 shows the absolute CBF values over time from fetal CI (baseline). Strategy alone did not impact any CBF parameter (p=0.62-0.81 mixed effects models) whilst time did (all p<0.0001). In all groups, an overall increase in mean and minimum, and a decrease in maximum and amplitude occurred following cord clamping. Mean, maximum and minimum waveform values were higher in the No-RM group compared with DynPEEP at the time of first inflations. All other timepoints and comparison of different ventilation strategies were comparable with no significant differences found in later timepoints. There was no difference in the CBF measures for the 14 cmH_2_O and 20 cmH_2_O DynPEEP levels (Online Supplementary Figure 1).

**Figure 1:**
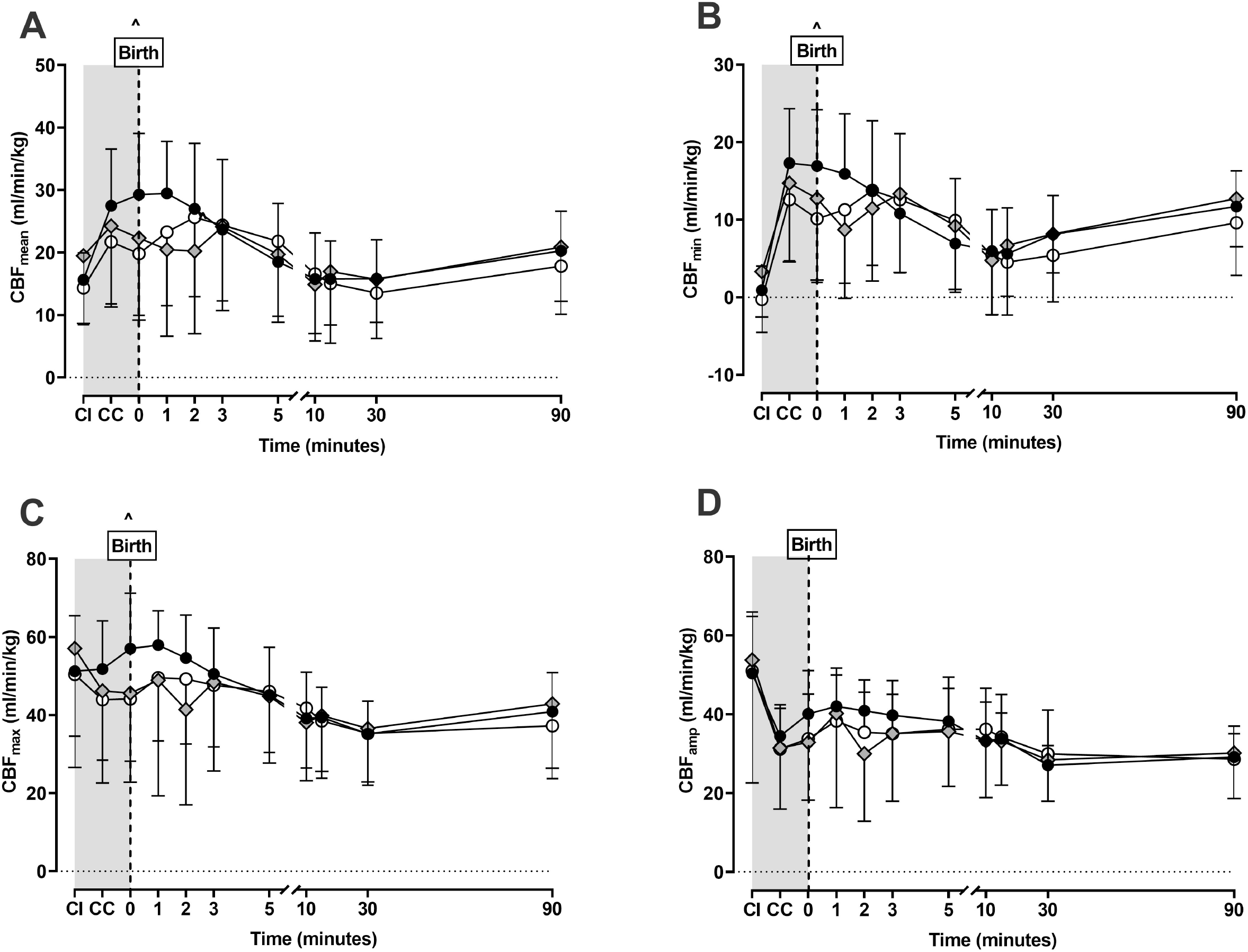
Absolute carotid blood flow (CBF) waveform measures including the mean (**A**; p<0.0001), minimum (**B**; p=0.0007), maximum (**C**; p<0.0012) and amplitude (**D**; p=0.15) during fetal measures with the cord intact (CI), following cord clamping (CC) before ventilation and 90 minutes of positive pressure ventilation in the No-RM (black circles), DynPEEP (open circles) and SI (grey diamond) recruitment groups. DynPEEP data represents pooled data for both the 14 and 20 cmH_2_O maximum PEEP. Fetal period without any ventilation shown in grey background. Birth; first 10s after commencing allocated recruitment strategy. All p values represent the overall p value using a mixed effects model (time and strategy combined). ^ No-RM vs DynPEEP p<0.05 Tukey post-tests. All data mean and standard deviation.

There was high inter-subject variability in all groups, including the initial CI value. To account for this, Figure 2 shows the change in CBF from fetal CI. Overall, time had the greatest impact on all CBF waveform measures (all p<0.0001 fixed effects), with mean CBF increasing after cord clamping, before decreasing to pre-birth levels by 10 minutes. Strategy alone only impacted mean CBF (p=0.049), whilst the combined impact of time and strategy was significant for mean (p<0.0001), minimum (p=0.0007) and maximum (p=0.0012) CBF data, but not amplitude (p=0.15). The change in mean and minimum CBF from fetal values was less during the first 3 minutes for the SI group. Thereafter the different ventilation strategies were comparable with no difference in later timepoints. In all groups, clamping of the umbilical cord caused the largest change in CBF (increase mean and minimum, decrease maximum and amplitude). Four lambs (3 DynPEEP; 7%, 1 SI; 7%) had a CBF that persisted at zero or negative values throughout the study (included in analysis).

**Figure 2.**
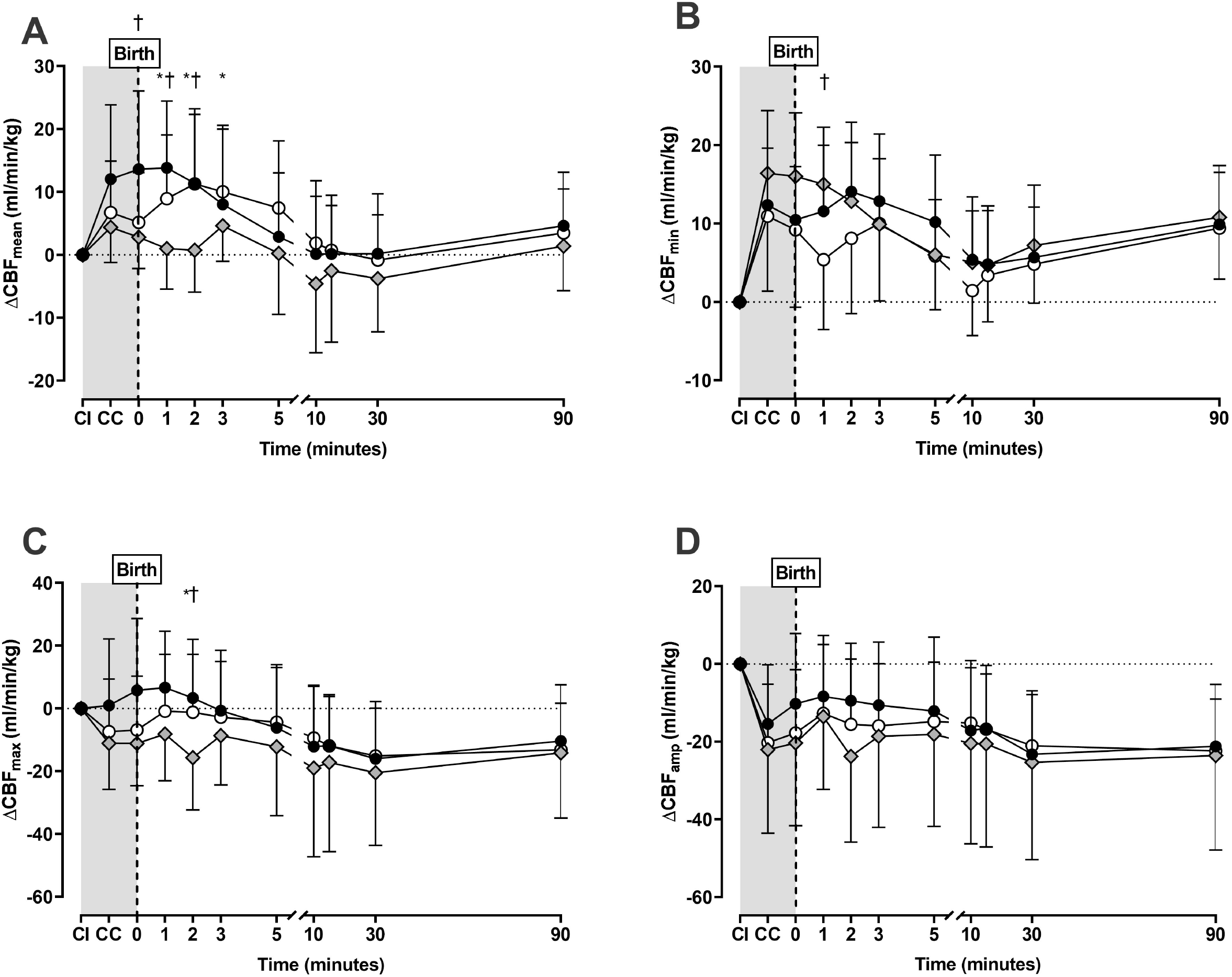
Change in mean (**A**; p<0.0001), minimum (**B**; p=0.0007), maximum (**C**; p=0.0012) and amplitude (**D**; p=0.15) of the carotid blood flow (CBF) waveform from fetal measures with the cord intact (CI), following cord clamping (CC) before ventilation and then 90 minutes of positive pressure ventilation in the No-RM (black circles), DynPEEP (open circles) and SI (grey diamond) recruitment groups. DynPEEP data represents pooled data for both the 14 and 20 cmH_2_O maximum PEEP. Fetal period without any ventilation shown in grey background. Birth; first 10s after commencing allocated recruitment strategy. All p values represent the overall p value using a mixed effects model (time and strategy combined). † SI vs No-RM, * SI vs DynPEEP p<0.05 Tukey post-tests. All data mean and standard deviation.

Figure 3 shows the absolute CAO content and carotid oxygen delivery over time. Irrespective of group, ventilation caused an increase in CAO content from fetal values (p<0.0001 for time). The combined impact of time and strategy was not statistically different (p=0.19). High variability in CAO content occurred in all groups at 5 minutes, with DynPEEP being lower, however, this was not significant. Carotid oxygen delivery increased similarly after birth for all groups (p<0.0001 for time and p=0.50 combined time and strategy). No difference was found in the CAO content and carotid oxygen delivery measures for the 14 cmH_2_O and 20 cmH_2_O maximum PEEP levels (Online Supplementary Figure 2).

**Figure 3.**
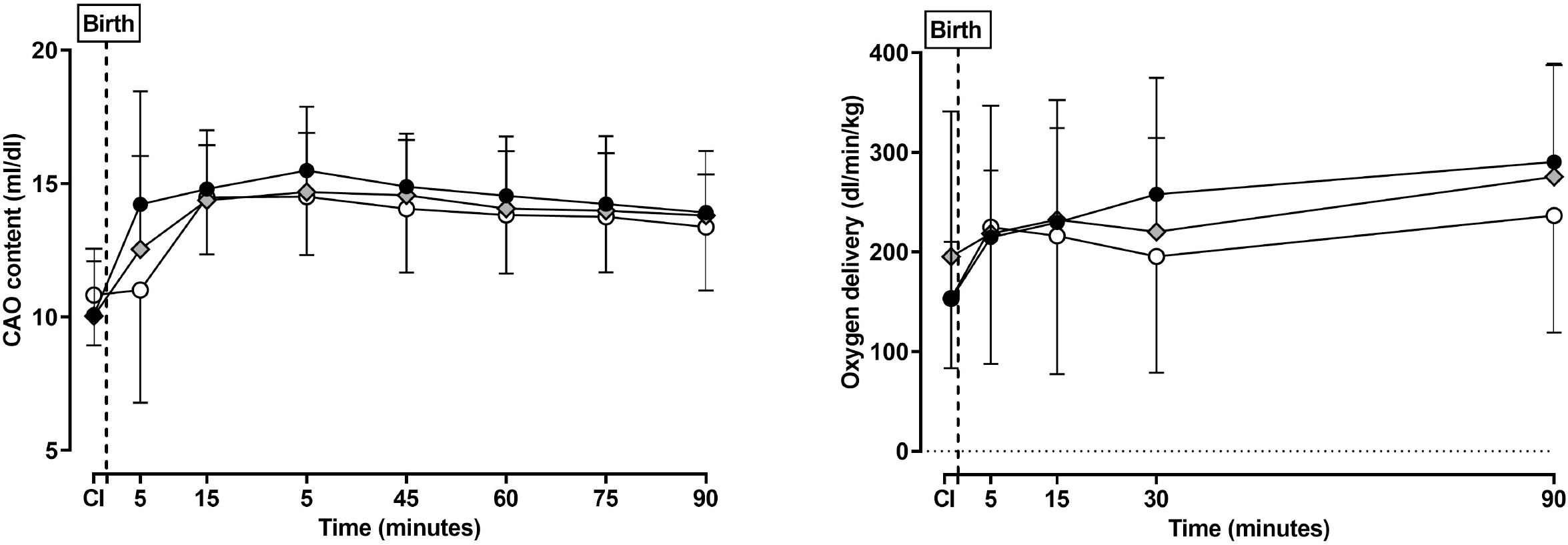
Absolute carotid arterial oxygen (CAO) content (**A; p=**0.21) and carotid oxygen delivery (**B;** p=0.50) in the fetal period (apnoeic cord intact (CI) and then at select timepoints during 90 minutes of positive pressure ventilation following the allocated 3 minute aeration strategy at birth for the No-RM (black circles), DynPEEP (open circles) and SI groups (grey diamonds). Birth; first 10s after commencing allocated recruitment strategy. All p values mixed effects model (time and strategy combined). All data mean and standard deviation.

## DISCUSSION

Supporting lung aeration at birth in preterm infants improves short-term respiratory outcomes.^1,2,18^ Despite the interest in different approaches in supporting aeration of the preterm lung at birth, systematic evaluation of cerebral haemodynamics has been limited.^11,14^ To our knowledge, our study is the first to assess the effect on cerebral haemodynamics of different PEEP strategies at birth. In general, we found that the aeration strategy did not cause short-term changes to CBF, CAO content or carotid oxygen delivery. Importantly, there was no significant difference in the use of transient PEEP levels between 8 and 20 cmH_2_O in the first 3 minutes of respiratory support. Fetal stability and cord clamping practices may play a bigger role in preterm cerebral haemodynamics at birth than the respiratory strategy.

In our study, the biggest change in CBF occurred after the umbilical cord was cut and before ventilation commenced, irrespective of group allocation. Cord clamping results in a wide range of vascular changes in preterm lambs, including an initial reduction in right ventricular output, initial increases in blood pressure over the initial 30s and stability of heart rate.^19^ This suggests that the effect of cord clamping itself is a prominent process that may influence CBF. CBF changes may have been different if the duration between cord clamping and ventilation was shorter, or ventilation commenced before cord clamping. ^20,21,22^ Although PEEP levels during ventilation prior to cord clamping have not been compared, our data suggests that the respiratory strategy is less likely to impact cerebral haemodynamics in the preterm infant than other factors such as cord care.

CBF is a commonly reported measure of cerebral haemodynamic physiology.^23,24^ We found that the transient increase in mean and minimum CBF at birth was less during the SI compared to No-RM and DynPEEP (initial 3 minutes). The high intra-thoracic pressure and, therefore, higher pulmonary blood flow and left ventricular output created by SI ventilation may explain this difference compared to the other strategies which uses tidal (cyclic) pressure changes. There is little agreement on the effects of SI on cerebral haemodynamics. A possible harmful effect of SI on cerebral oxygenation (measured with Near-InfraRed Spectroscopy) was found in a small study of preterm infants.^25^ Whilst both an increased risk of cerebrovascular injury, CBF and carotid oxygen delivery,^17^ and a reduction in CBF compared to non-SI interventions have been reported in preterm lambs.^11^ These conflicting results may reflect the small sample sizes in each study (approximately 6/group). This is particularly relevant given the high-level of inter-subject variability that we observed in our larger SI sample of 15, highlighting the importance of appropriately powered preclinical studies. We have also previously reported a high variability in the time and volume response needed to appropriately aerate the lung using a SI.^6,26,27^ There is no accepted definition of an ‘optimal’ SI approach, with pre-defined, and fixed, time being the most common. ^7,28,29^ The use of a fixed duration SI is based on the false assumption that all recipients lung mechanics will respond the same.^6,26,27^ We have previously shown that titrating the SI duration to the aeration response in real-time results in better respiratory outcomes than the 30s SI most commonly used in lamb studies.^6^ In our earlier studies, a 30s SI could be too short or unnecessarily excessive in terms of lung aeration.^6^ This may explain the mixed results in preclinical and clinical preterm SI studies, especially the unexplained higher early mortality in the SI group in the Sustained Aeration of Infant Lungs (SAIL) trial.^30^ Whilst titrating the SI duration to response is not used clinically, in a preclinical study it is arguably preferable as it standardises the intervention across all lambs irrespective of the lung mechanics. Even with a titrated strategy to aeration, a SI provides no benefit in lung aeration and increases early injury markers compared to DynPEEP and No-RM.^2^ The clinical implications of the different CBF patterns found in this study are unknown and cannot be used to recommend one respiratory support strategy over another.

CAO content provides a measure to assess the availability of oxygen in the blood. CAO content increased from fetal values for all ventilation strategies, indicating that each group supported cerebral oxygenation similarly and successfully. Interestingly, the DynPEEP group did not show an initial increase in CAO content from the fetal baseline until 15 minutes, compared to 5 minutes in the SI and no-RM groups. The exact cause for this is unclear, and we can only speculate on the reasons. We have previously reported a lower dynamic respiratory system compliance prior to ventilation in this group of DynPEEP lambs at birth,^2^ suggesting a possible fetal difference in the groups. The high variability indicates that the likely reason is a degree of transient hypoxia or asphyxia following cord clamping (weigh and moving), which is known to occur with as little as 45s delay in ventilation.^15^ Importantly, the CAO content was similar in all groups by 10 minutes suggesting that any effect is transient. Irrespectively, these findings indicate that the higher PEEP at birth may impact cerebral oxygen delivery. Future well powered studies with more arterial sampling intervals in the first 10 minutes are required. As we cannot conclude that there is not a potential impact on CAO content, clinical trials of DynPEEP, such as Positive End-Expiratory Pressure Levels during Resuscitation of Preterm Infants at Birth (the POLAR trial),^31^ should include cerebral complications in their analysis.

Carotid oxygen delivery, as a function of blood flow and oxygen content, is a thorough and accurate assessment of carotid haemodynamics. We found that the carotid oxygen delivery similarly increased from fetal levels in all groups, consistent with other comparisons between SI and No-RM in preterm lambs.^11,17,32^ Some preterm lamb studies have indicated a significant increase in oxygen delivery in the No-RM group compared to the SI groups, when ventilated with 100% oxygen.^11^ Whilst this may indicate that the level of oxygenation influences cerebral haemodynamics, the lack of difference in our groups at 5 minutes (oxygenated in a standardised 30% inspired oxygen) supports our assertion that achieving lung aeration rather than the means of achieving it is the critical determinant of cerebral haemodynamics.^2^

Despite the lung protective benefits of DynPEEP at birth in preterm lambs,^2,4^ this strategy may result in detrimental cardiovascular impacts. Impairment of pulmonary blood flow has been reported in a previous study of lambs managed with a DynPEEP approach between 4 and 10 cmH2O after respiratory transition.^12,13^ It should be noted that lung aeration was not measured in the studies of Polglase and co-workers, limiting any interpretation of the role of volume state (overdistension or recruitment). Our study has been the first to assess the use of DynPEEP on cerebral haemodynamics during the respiratory transition, and we have previously reported that overdistension was not a feature in our lambs.^2^ Thus, it is interesting that PEEP levels (between 8 and 20 cmH_2_O) did not significantly alter CBF or cerebral oxygen delivery. An important aspect of these PEEP strategies is the transient high PEEP exposure whilst the lung is clearing fetal lung fluid. Current clinical approaches to DynPEEP also constrain PEEP increases to clinical need, based on oxygen and heart rate response,^31,33^ rather than pre-fixed PEEP values which should further limit any potential risk of inadvertent high intrathoracic pressure on haemodynamics.

A redirection of CBF from the brain was noted in four lambs, most likely during diastole. It is possible that a redirection of blood flow to the ductal and pulmonary arteries caused this.^15^ This cannot be confirmed as pulmonary or ductal blood flow was not measured in our study. As DynPEEP studies in infants are now being conducted, understanding the physiological impact of PEEP levels on pulmonary and ductal blood flow is needed.

This study has limitations. Data was sourced from a large biobank of inter-linked preterm lamb studies primarily assessing lung injury resulting in unbalanced group sizes. Cerebral haemodynamics were not the primary outcome of these studies, leading to some lambs being excluded due to unreliable CBF data (but not clinical outcomes). In addition, we found little difference between the three ventilation strategies. Our study was not designed as a non-inferiority or equivalence trial and conclusions regarding cardiovascular and cerebrovascular safety should be made with caution. We studied anaesthetised apnoeic and intubated lambs, a common preclinical model^2,11,13,17,26^ but not representative of clinical practice. In our study, cerebral flow dynamics were assessed from a single carotid artery, with occlusion of the other. Whilst this is common practice in lamb studies,^22,34^ it is not possible in humans. Ultrasound measures or Near-InfraRed Spectroscopy have been used in humans, including preterm infants, to assess CBF and cerebral oxygenation,^35,36, 37^ and may have aided translatability. Extrapolating unilateral carotid blood flow measures to global cerebral haemodynamics requires caution, as unilateral occlusion may result in changes to the flow dynamics of the contralateral side.^24^ Further, our measures should not be assumed to represent the cerebral microvascular bed, with a recent preterm lamb pilot study suggesting large and small cerebral vessel behaviour is not always uniform.^14^ Finally, we did not measure cerebral injury as an outcome.

## Conclusion

There is a paucity of information on the cerebral haemodynamic effects of respiratory strategy at birth in preterm infants, despite the clear risk of injury in this population. Reassuringly, respiratory strategy did not impact cerebral haemodynamics, suggesting that clinicians can focus on strategies that optimise cardiorespiratory outcomes during the transition to air breathing.

## Supporting information

ARRIVE Criteria Check List

## Acknowledgments

The authors acknowledge Regina Oakley for assistance in lamb management and preparation of some of the gas exchange and lung mechanics data, and Sarah White and Rebecca Sutton for assistance in preparation of the ewes, and Jean Hellstern for proofreading the manuscript.

## Author contributions

DGT and PP-F developed the concept and designed the experiment. DGT, PP-F, EJP, KEMcC, MS were involved in lamb experimental work. SD, KRK, DS performed the carotid and cerebral data analysis. All authors participated in data interpretation. DGT supervised all aspects of the study and subsequent data analysis. SD wrote the first draft and all authors contributed to redrafting the manuscript.

## Ethics Approval and Reporting

The study was approved by the Animal Ethics Committee of the Murdoch Children’s Research Institute, Melbourne (MCRI), Australia (A780, A790, A822) in accordance with National Health and Medical Research Council guidelines and is reported as per the ARRIVE guidelines.

## Data Sharing

All data, including raw data used for all figures and analysis, is available upon request to the corresponding author from three months following article publication to researchers who provide a methodologically sound proposal, with approval by an independent review committee (“learned intermediary”). Proposals should be directed to to gain access. Data requestors will need to sign a data access or material transfer agreement approved by MCRI.

