## Supplementary Figure 1 and 2 and Table 1 for "Aeration Strategy at Birth Does Not Impact Carotid Haemodynamics in Preterm Lambs"

**Aeration Strategy at Birth Does Not Impact Carotid Blood Flow and Oxygen Delivery in Preterm Lambs**

**Online Data Supplement**

**Online Table 1.** Dynamic 14 cmH_2_O and 20 cmH_2_O PEEP Group Lamb Characteristics

**Online Figure 1.** Absolute carotid blood flow (CBF) waveform measures for the 14 cmH_2_O and 20 cmH_2_O maximum PEEP Dynamic strategies.

**Online Figure 2.** Absolute carotid artery oxygen (CAO) content and carotid oxygen delivery for the 14 cmH_2_O or 20 cmH_2_O maximum PEEP Dynamic strategies.

**Online Table 1.** Dynamic 14 cmH_2_O and 20 cmH_2_O PEEP Group Lamb Characteristics

| **Dynamic PEEP Group** | **n** | **GA (d)** | **Weight (kg)** | **Gender**  **(F:M)** | **Parity**  **(S:T)** | **Fetal Fluid (ml/kg)** | **Static C_RS_ (ml/kg/cmH_2_O)** | **Fetal Arterial Blood Gas** | | | | **Cerebral Blood Flow (ml/kg/min)** | | | |
| --- | --- | --- | --- | --- | --- | --- | --- | --- | --- | --- | --- | --- | --- | --- | --- |
|  |  |  |  |  |  |  |  | **pH** | **PaCO_2_**  **(mmHg)** | **PaO_2_**  **(mmHg)** | **BiC**  **(mmol)** | **Height** | **Min** | **Mean** | **Max** |
| **Low** | 14 | 125.5 (1.0) | 3.52 (0.32) | 8:6 | 0:14 | 16.6 (5.0) | 1.18 (0.22) | 7.36 (0.07) | 47.0 (7.4) | 27.9 (3.0) | 24.2 (2.7) | 52.3 (14.2) | -0.6 (5.1) | 15.0 (5.7) | 50.7 (14.9) |
| **High** | 27 | 125.2 (1.0) | 3.33 (0.32) | 18:9 | 3:24 | 17.8 (5.2) | 1.22 (0.19) | 7.34 (0.06) | 47.5 (6.9) | 27.3 (6.3) | 23.0 (2.6) | 50.3 (17.9) | -0.1 (3.8) | 14.0 (6.1) | 50.3 (16.6) |

**Abbreviations:** GA; gestational age, F; female, M; male, S; singleton, T; multiparity, C_RS_; compliance, PaCO_2_; partial arterial pressure of carbon dioxide, PaO_2_; partial arterial pressure of oxygen, Bic; bicarbonate. All data mean (SD) or ratio. All p values one-way ANOVA or chi-test as appropriate.

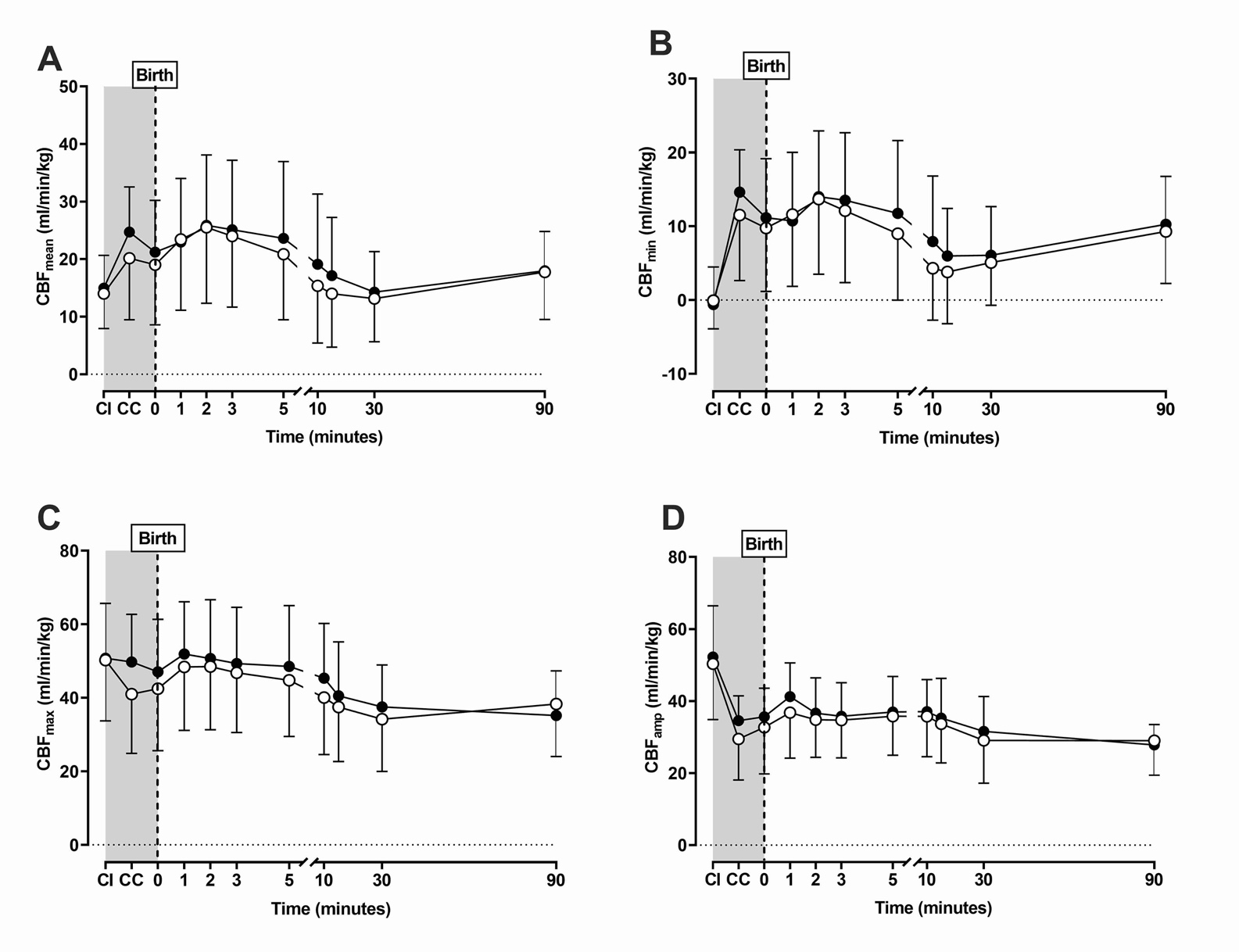
**Online Figure 1.** **Absolute carotid blood flow (CBF) waveform measures for the 14 cmH_2_O and 20 cmH_2_O maximum PEEP Dynamic strategies.**

**Online Figure 1:** Absolute carotid blood flow (CBF) waveform measures including the mean (**A**; p=0.9479), minimum (**B**; p=0.9043), maximum (**C**; p=0.6684) and amplitude (**D**; p=0.9024) in the 14 cmH_2_O (black circles) or 20 cmH_2_O (white circles) maximum PEEP Dynamic strategies. All p values overall mixed effects model (time and strategy combined). There was no difference in the groups at each time point (all p>0.05; Tukey post-tests). Fetal period without any ventilation shown in grey background. CI; cord intact, CC; post-cord clamping, Birth; first 10s after commencing allocated recruitment strategy. All data mean and standard deviation.

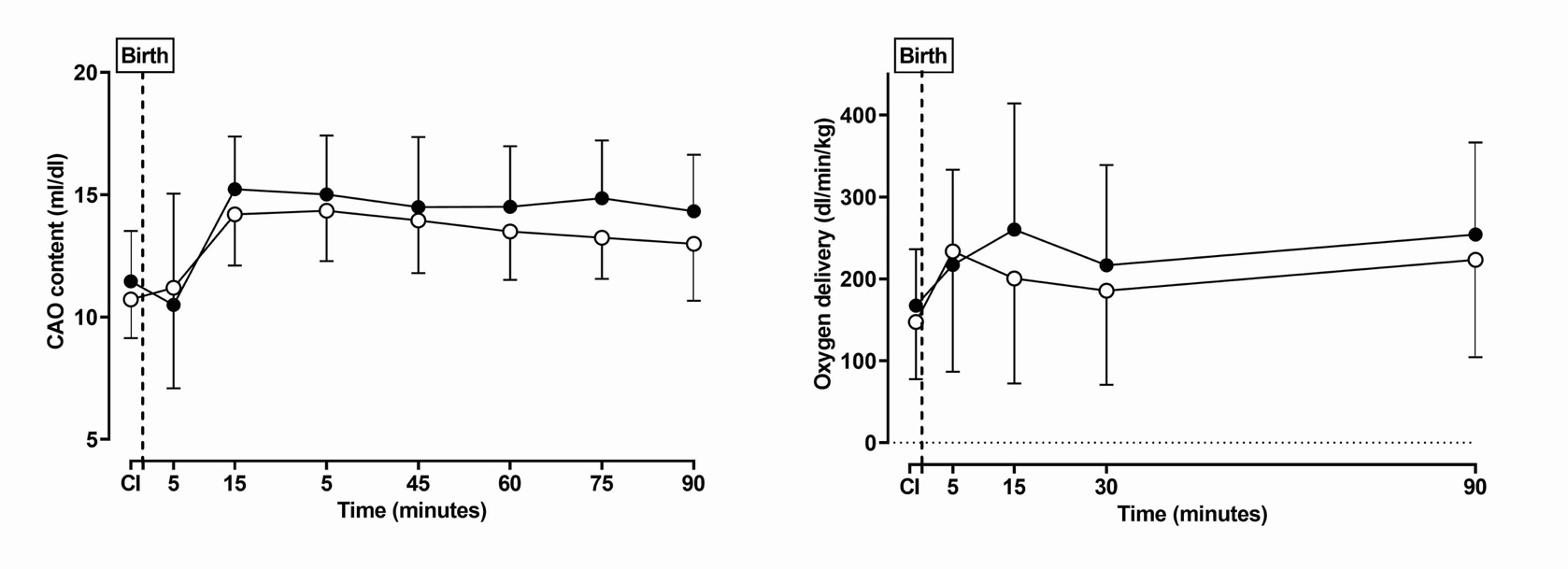
**Online Figure 2. Absolute carotid blood flow, carotid artery oxygen (CAO) content and carotid oxygen delivery for the 14 cmH_2_O or 20 cmH_2_O maximum PEEP Dynamic strategies.**

**Online Figure 2:** Absolute carotid arterial oxygen (CAO) content (**A;** p=0.2712) and carotid oxygen delivery (**B;** p=0.2800) in the 14 cmH_2_O (black circles) or 20 cmH_2_O (white circles) maximum PEEP Dynamic strategies. P values calculated with an overall mixed effects model (time and strategy combined). There was no difference in the groups at each time point (all p>0.05; Tukey post-tests). CI; cord intact, CC; post-cord clamping, Birth; first 10s after commencing allocated recruitment strategy. All data mean and standard deviation.
